# Computational Models of Age-Associated Cognitive Slowing

**DOI:** 10.1101/2024.06.24.600545

**Authors:** Sanya Ahmed, William W Lytton, Terrence Stewart, Howard Crystal

## Abstract

**Background:** Cognitive slowing accompanies normal aging, yet understanding of the mechanisms of slowing is limited at the network and neuronal level. Relating the pathophysiological factors responsible for cognitive slowing, and interpreting its relationship to working memory, requires multiscale computer modeling.

**Objective:** The aim of this research is to explore multiple mechanisms of cognitive slowing using computational modeling of the cortex to link neuronal activity with cognitive content.

**Method:** We developed multiscale computer models of a simple cognitive task - Condition 1 of the Stroop recognition task - using the Nengo system, a cognitive simulation environment with a semantic pointer architecture developed to model cognitive tasks using spiking neural networks. We explored how changes associated with aging such as increased input noise, axonal loss, neuronal loss, and feedback would affect the function of the models.

**Results:** Axonal loss and increased input noise produced profound slowing. High levels of neuronal loss severely impaired memory and paradoxically decreased slowing via the ability to respond more quickly by “releasing” a prior memory. Increased feedback improved memory at the cost of increased slowing.

**Conclusion:** Our simulations suggest that significant slowing could be caused by white matter loss (axonal loss) or input signal degradation (which could be caused by visual or other afferent system worsening). As neuronal loss markedly decreased the duration of working memory, we propose that physiological feedback is increased to preserve working memory at the cost of further cognitive slowing.

## Introduction

Slowing on cognitive tasks is a part of normal aging. For example, the average score for 70 years-old subjects on the timed digit-symbol test was 70% of the average score for 20 year-olds on the same test, with multivariate analysis demonstrating that age accounted for 86% of the variance (Hoyer et al., 2014). The more complex the cognitive task, the more pronounced the difference of speed between age groups. While the average speed on simple reaction time tasks declined 9% from 20 to 60 years of age, the average speed on choice reaction time tests declined between 17% to 38% depending on the test (Woods et al., 2015). Times on the Stroop interference task show order of magnitude effects: from 89 ms at age 25 to 334 ms at 63 to 813 ms at 78 (Wolf et al., 2014).

Cognitive slowing accounts for much of age-associated cognitive decline across multiple domains (Cerella and Hale, 1994; Birren and Fisher, 1995; Cabeza et al., 2018; Salthouse, 1996a, 1996b, 2016, 2017; Ebaid, Crewther et al., 2017). Slowing is further exacerbated in specific dementia syndromes where it can progress to cognitive breakdown, thereby producing functional impairment. Although cognitive slowing is a feature of all types of dementia, ‘subcortical’ dementias such as Parkinsonian and Parksinson-plus dementias, normal pressure hydrocephalus, and some forms of vascular dementia present notably severe slowing (Huber, Shuttleworth et al., 1986; Cummings, 1986; Pillon, Dubois et al., 1991; Bonelli and Cummings, 2008).

The neural substrate accounting for slowing is unknown, and it is likely that multiple factors contribute, with distinct factors dominating in different forms of age-related and dementia-related dysfunction. These factors include: 1) increased input signal noise or decreased signal strength due to reduced efficiency of sensory organs: eyes, ears, touch; 2) limiting the transmission of action potentials via axonal loss, resulting in increased interneuronal synaptic delays (Turken et al., 2008; Kerchner et al., 2012; Legon et al., 2016); and 3) neuronal loss, thought to produce reduced drive along particular neural pathways. Understanding and differentiating the neural substrates of cognitive slowing would permit development of biomarkers to differentiate particular types of dementia in the preclinical stage and may provide clues to therapeutic strategies to mitigate cognitive slowing.

In the first condition of the original ‘paper and pencil’ Stroop, 100 samples of red, green, or blue are aligned 10 per row on 10 rows, and the subject is timed while naming all 100 colors (Scarpina and Tagini, 2017). With the computerized test, the subject presses a key corresponding to red, green, or blue after the new stimulus is presented (Figure 1). In this example, which we use for the computer model, the color red is presented after a one second delay followed by blue at two seconds and then green at three seconds. *In vivo,* the subject would press a key corresponding to the color as quickly as possible after the color changes. Reaction time (RT) is the delay from color presentation to key-press time. RT will be preceded by *time to registration* (TTR) of the color by a region or several regions of cortex. *In silico*, we simulate one cortical area and term the TTR as the time needed to reach a threshold representing recognition.

**Figure 1.**
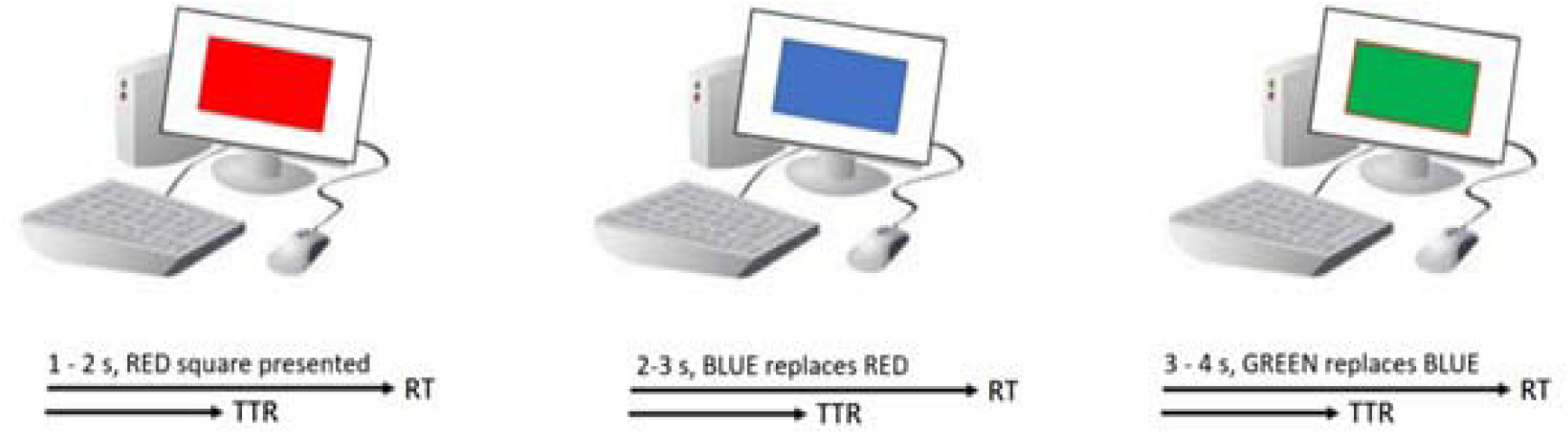
Condition 1 of the Stroop test: a color stimulus is presented and the subject presses a key to identify the color. The time between presentation of the stimulus and recognition (internal representation) is termed the *time to registration* (TTR) which precedes the time to key-press (the reaction time (RT)).

Our basic computer simulation of Stroop Condition 1 used the Nengo simulation system to explore: 1) input noise; 2) axonal loss; 3) neuronal loss; and 4) recurrent excitation strength. We found that parameter changes produced dysfunction indicated by increased TTR, decreased working memory duration, or both. Our results demonstrated that a combined increase in all parameters within a normal range associated with aging would be expected to yield both cognitive slowing and memory impairment. Increasing the degree of recurrent excitation would compensate for memory loss at the cost of further slowing.

## Methods

We identify 1) The Neuroengineering Framework; 2) Equations used for encoding of the input signal into neuronal activity and decoding of neuronal activity back to signals; 3) Vector and dot-product representations of decoded spiking activity from the computer model; 4) Definition of time to registration (TTR) and memory duration; 5) Representation of input noise; 6) Simulation of axons; 7) Modeling neuronal ablation; 8) Parameter space, predictions, and hypotheses; and 9) Code and data repository.

### 1) The Neuroengineering Framework

As described in Fig 1 of Bekolay et al., the Neural Engineering Framework (NEF) provides encoding, internal continuing representation, and decoding to provide a recognition loop. In the complex systems developed by the originators of NEF, this sequence is made to represent a brain area, and brain areas including striatum, thalamus, and cortex are then connected in an anatomically plausible fashion. In our case, we are using a single ensemble to do a simple encoding/store/decoding task. The inputs are three sets of labeled lines that represent the three colors to be registered. Each neuron in the ensemble has its own randomly assigned tuning curve. Comparable to the tuning curves of cone cells in the retina, individual neurons will respond most strongly to one color but respond in varying degrees to the other colors as well.

The strength of total input into a particular neuron will translate into probability of firing on a moment-by-moment basis. By the representation principle, the spiking activity of a neural population can be decoded to recover the original input signal, or some transformation of that input signal. First, the firing pattern is filtered with a decaying exponential filter. The filtered activity is then summed together with a set of weights that approximates the input signal and the cosine of the input signal. By the transformation principle, populations of neurons can send signals to another population by decoding the desired function from the first population and then encoding the decoded estimate into the second population. These two steps can be combined into a single step by calculating a set of weights that describe the strength of the connections between each neuron in the first population and each neuron in the second population. A neurally implemented dynamical system has negative feedback across its two dimensions, resulting in a harmonic oscillator. By the dynamics principle, signals being represented by a population of neurons can be thought of as state variables in a dynamical system.

Because these computer modeling techniques may be unfamiliar to readers trained in behavioral neurology/neuropsychology, we provide here a brief introduction to the Neuroengineering Framework (NEF) using the *Nengo* language found on www.nengo.ai (Bekolay, Bergstra et al., 2014; Eliasmith et al., 2012; Eliasmith, 2013). In NEF, digital input is encoded as the spiking activity of simulated integrate-and-fire neurons which are grouped into neuronal ensembles. At the input level, neurons have different tuning curves so that the group spans the representational space, which in this case is color. Neurons within an ensemble are fully interconnected: they all receive the same input and project to the same output. Inhibitory neurons are not explicitly modeled but negative synaptic weights are used. Maximal neuronal firing rates are set to 200 Hz. Groups of cells are considered vectors for the purpose of encoding and decoding and are encoded into spiking activity of simulated neurons. The output of spiking activity is subsequently decoded back into vectors (Eliasmith et al., 2012; Eliasmith 2013; Bekolay et al., 2014).

The models utilize the ‘semantic pointer architecture’ of the NEF, summarizing the spiking activity of thousands of neurons from one or more ensembles, comparable to the use of group activity in functional magnetic resonance imaging (fMRI) studies of semantic memory. One way to summarize that spiking activity is with a vector of many elements, each of which represents a different set of neurons within the cortical slab. There are multiple data reduction methods which can be implemented to summarize neuronal activity. Each method must be evaluated for information loss under different conditions and biological plausibility. The rationale and methodology of the method used in Nengo has been discussed previously.

We modeled a single ‘slab’ of the cortex that is similar in its information processing function to visual area V4 in the extrastriate visual cortex. The model consisted of 6400 simulated integrate-and-fire neurons. The neurons were organized into eight ensembles each containing 800 neurons. The spiking activity of each ensemble was summarized as a vector containing 16 elements. The activity of the eight ensembles could thus be summarized by a vector containing 128 elements. Control simulations showed that comparable results were obtained with other ensemble architectures as well. We grossly conceive of the neuronal ensemble as organized similar to microcolumns. In the model, neurons within each ensemble are fully interconnected, including self-connects. There are no connections across ensembles (Figure 2).

**Figure 2.**
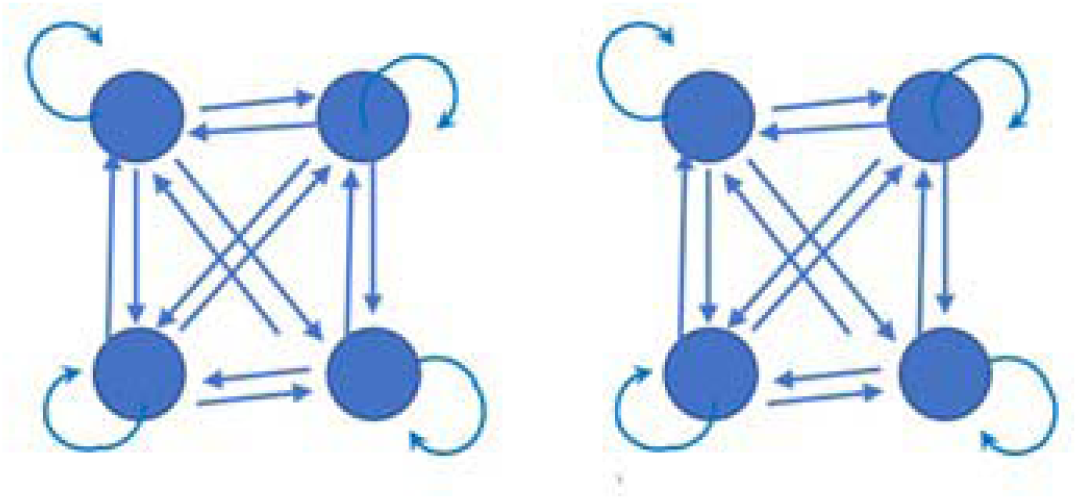
Schematic of two ensembles, each containing four neurons represented by circles. The full model contains eight such ensembles of 800 neurons each.

In our computer models, we are representing the input into the modeled cortical slab as a 128-element vector. Simulated responses to the colors red, blue, and green are represented by different arbitrarily generated vectors which overlap by 10%, respectively named ‘RED’, ‘BLUE’, and ‘GREEN’. Although we envision a process similar to the encoding steps happening within biological brains, we do not conceive of the decoding steps as brain-like; However, the decoding steps are necessary to assess the accuracy and dynamics of the encoding steps within the simulation.

### 2) Equations used for encoding of the input signal into neuronal activity and decoding of neuronal activity back to signals

The input vector **x** is transformed into spiking neuronal activity via Equation 1.

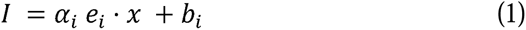

The input current is calculated by multiplying the input vector by the gain (a_i_) and the basis vector (e_i_) and then adding the bias (b_i_). The input current is then transformed into spiking activity within a leaky integrate-and-fire neuron.

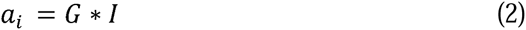

Where G is a neural non-linearity. In this work, we modeled a leaky-integrate-and-fire (LIF) neuron with a time constant of 20 ms and an absolute refractory period of 2 ms.

Following this encoding process, we provide a decoding so that the system can identify the color perception. The task is to identify a subset of the set of decoding weights **d** that when multiplied by neuronal activity (**a**) yields the output vector (**x**).

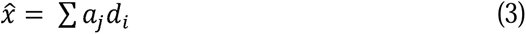

To find **d**, the NEF first computes the sum of the simulated neuronal activity over a random sample of x values, represented by the variable Y:

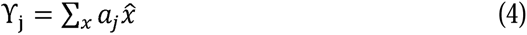

The NEF then calculates the matrix formed by the vector products of encoding and decoding weights, denoted I’:

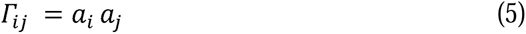

Finally, **d** is computed by taking the inverse of I’:

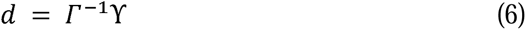

Unless explicitly specified otherwise, default software values were used for all simulations.

### 3) Vector and dot-product representations of decoded spiking activity from the computer model

In order to determine how accurately the spiking activity across ensembles represents the signals input into the models, the activity of 6400 neurons was decoded into a 128-element output vector (Figure 3). The decoded output changes for each element every ms. A plot of all 128 lines representing the ms by ms decoded output is difficult to interpret. By plotting fewer elements, the figure is tractable, however, its relationship to color representation is uninterpretable (Figure 3a). Clarity is achieved by computing the ms by ms dot products between the 128-element output and the predefined 128-element input vectors representing ‘RED’, ‘BLUE’, and ‘GREEN’ to create an array of one-dimensional scalar values (Figure 3b). The model quickly represents ‘RED’ in the first second. When ‘BLUE’ is input at two seconds, the model takes time for the representation of ‘RED’ to descend and the representation of ‘BLUE’ to ascend.

**Figure 3.**
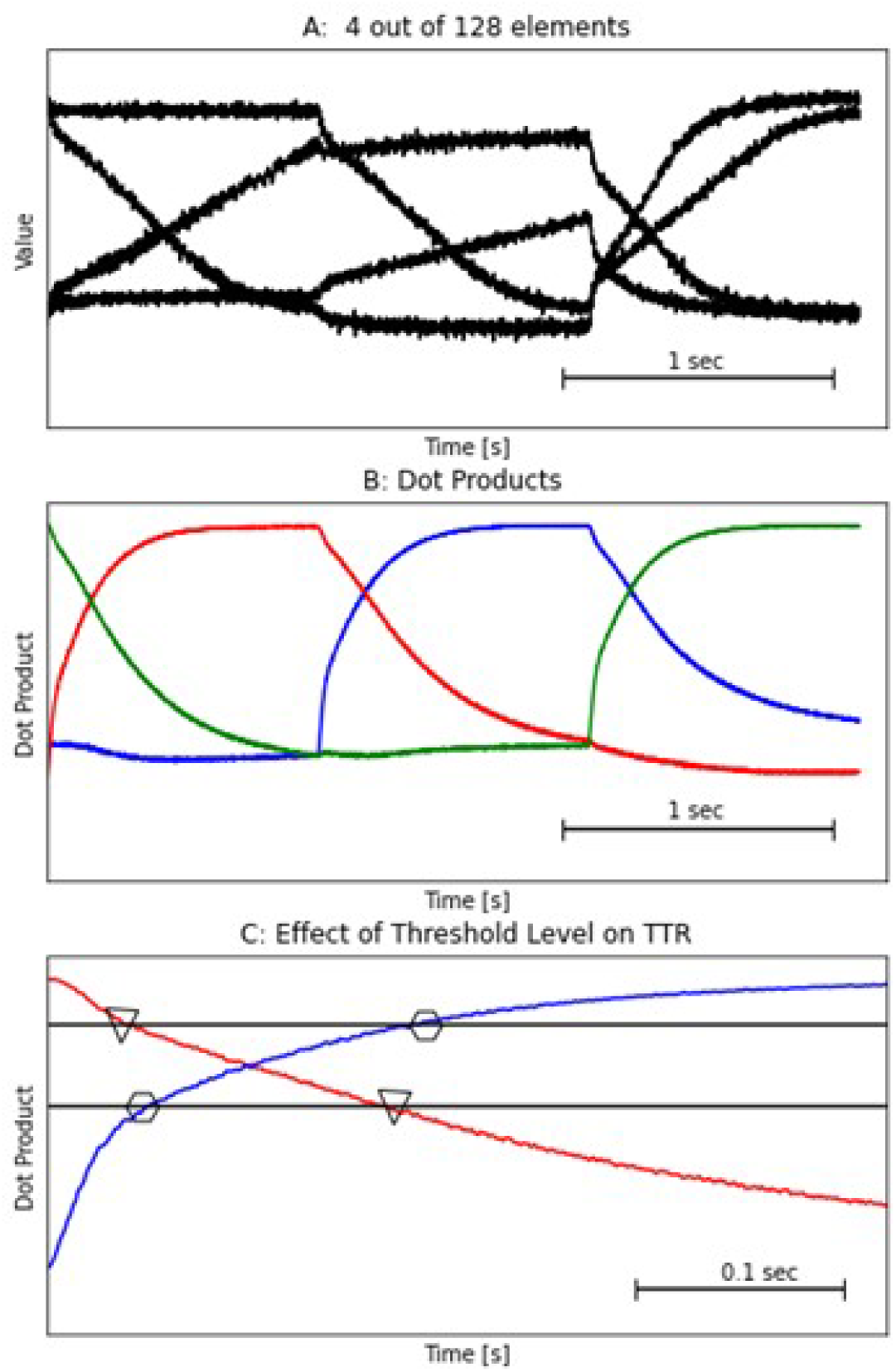
Decoded outputs of the spiking activity across all eight ensembles, with each element in the vector plotted as a function of time. A: Four arbitrary elements out of the total 128. B: The ms by ms dot products of these vectors, yielding scalar values with less deviation. C: The dot products in detail over a 0.5 second interval of time during which the ascending ‘BLUE’ and descending ‘RED’ vectors intersect, superimposed by the high (0.8) and low (0.5) threshold levels.

### 4) Definition of time to registration (TTR) and memory duration

We defined TTR as the time required for the representation of the ascending input to cross a threshold of 0.8. The time for the descending, incorrect response to fall below the threshold is defined as the duration of working memory.

### 5) Representation of input noise

Noise and randomness are built into NEF simulations at multiple levels (Bekolay et al, 2014). Input noise was modeled as an arbitrarily defined 128-element vector similar in concept to ‘RED’, ‘GREEN’, or ‘BLUE.’ The signal strength of the ‘RED’, ‘GREEN’, and ‘BLUE’ input vectors were varied from 100% to 40%, and the strength of an additional vector representing the input noise, termed ‘NOISE’, was varied from 0% to 60% accordingly. Input signals consisted of a combination of some fraction of correct signal (ranging from 0.4 to 1.0) and some fraction of ‘NOISE’ (ranging from 0.0 to 0.6).

### 6) Simulation of axons

In order to accurately simulate aging and disease, in which axons can be lost altogether, we modeled axonal loss by decreasing the projection of input signal into the ensemble by a proportion of up to 20%.

### 7) Modeling neuronal ablation

Ablation was performed by setting the encoder and gain associated with a neuron to zero. Neuronal ablation was performed up to 20% of the total neuron count.

### 8) Exploring parameter space, predictions, and hypotheses

TTR measurements were obtained for four varied parameters in the simulations: 1) the percentage of noise in the input signal; 2) the percentage of intact axons; 3) the percentage of neuronal ablation; and 4) the degree of recurrent excitation.

We developed four initial hypotheses regarding the behavior of the simulation in response to varied parameters: 1) decreased signal strength or increased input noise will increase the TTR; 2) axonal loss will increase the TTR; 3) neuronal ablation will increase the TTR; and 4) A compensatory increase in the gain of recurrent synapse will improve memory duration at the cost of TTR slowing.

### 9) Code and data repository

Code and simulated data will be available at modeldb.yale.edu.

## Results

### 1) The Effects of Threshold Level on TTR

Threshold levels above and below the intersection of 0.65 where the descending representation of the previous, no longer correct color (red), crossed with the ascending representation of the current, correct color (blue) affected the TTR (Figure 3c). At a high threshold of 0.8, representation of ‘RED’, the incorrect response, persisted for 50 ms. Between 50 and 180 ms, neither the representation of ‘RED’ nor ‘BLUE’ was above the threshold: we call this interval ‘No Response’. After 180 ms, the representation of ‘BLUE’ was above the threshold (Figure 3c). We defined this unambiguous response, where one color is above threshold, as the TTR.

A threshold value below the intersection of the ascending, correct, and descending, incorrect, color led to an ambiguous response where both colors were represented. With a low threshold of 0.5 (below the intersection), the incorrect response persisted until the representation of ‘RED’ descended below threshold at 180 ms. The correct response occurred when the representation of ‘BLUE’ crossed the threshold at 45 ms. Thus, between 45 and 180 ms, the response was ambiguous as representations of both ‘RED’ and ‘BLUE’ were above threshold. The simulations presented in this work used the higher threshold of 0.8 which avoids ambiguous responses.

### 2) Increased input noise caused TTR slowing

Input noise increases with aging due to deterioration of the sensory apparatus. In the visual domain, the visual stimulus may degrade due to cataracts, glaucoma, retinal degeneration, etc. Similarly, signals from the auditory domain will degrade due to both inner ear and middle ear changes. In our model, added input noise produced modest slowing up to 30%, with profound slowing seen at 60% (Figure 4). Beyond 60% input noise, representation was not possible as the ability to recognize the input stimulus failed.

**Figure 4.**
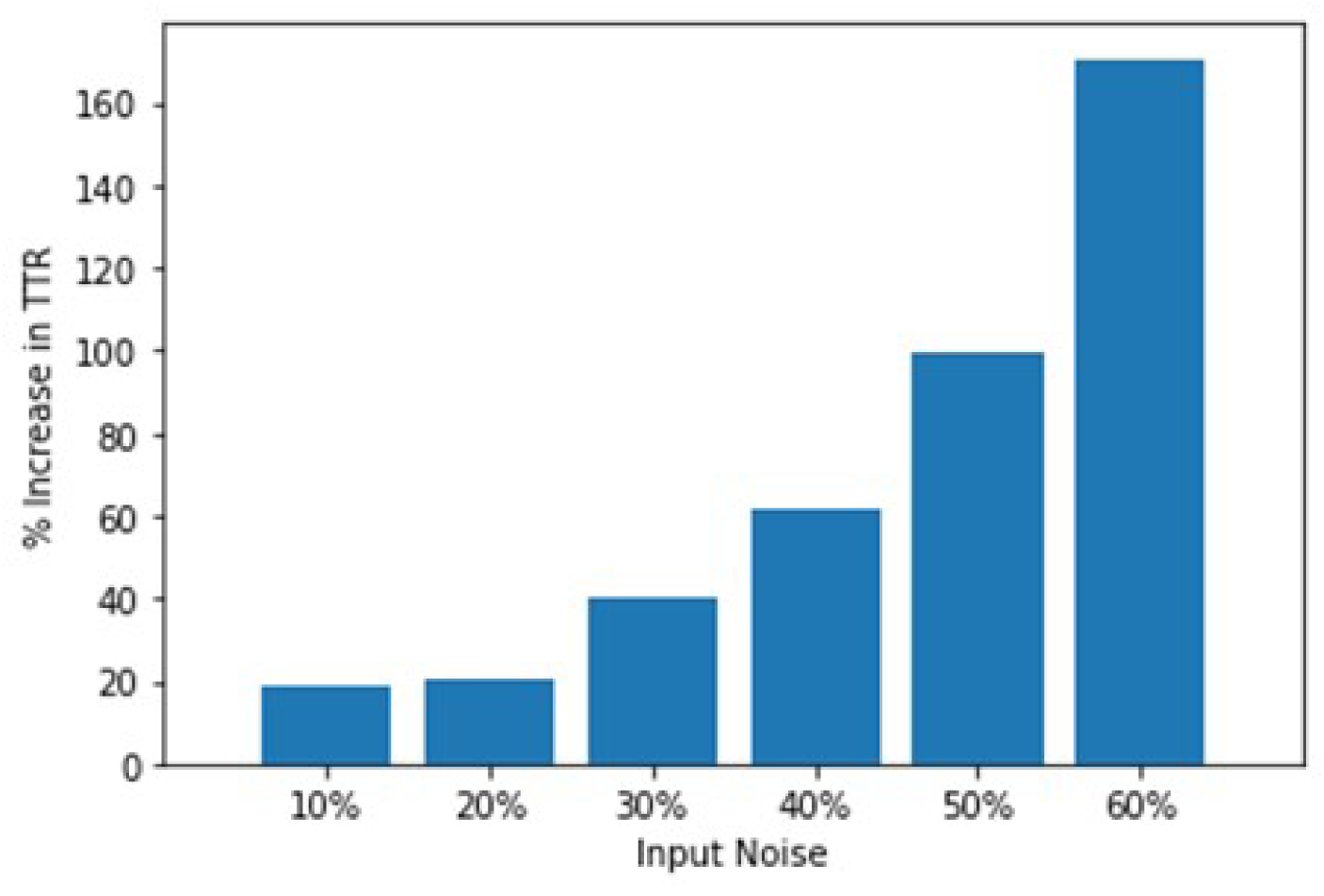
Added input noise increased the TTR.

### 3) Axonal loss caused TTR slowing

Axonal damage is among the first degenerative changes in the aging brain, leading to a loss of synaptic input which disrupts the spatiotemporal interactions between ensembles. (Anhoque et al., 2013; Salvadores et al., 2017; Wolf and Koch, 2016). TTR with 80% of the original number of axons was notably slower than when axons were fully intact (Figure 5). Abundant firing was needed for inputs to cross the TTR threshold, a process prolonged by fewer synapses in axonal loss.

**Figure 5.**
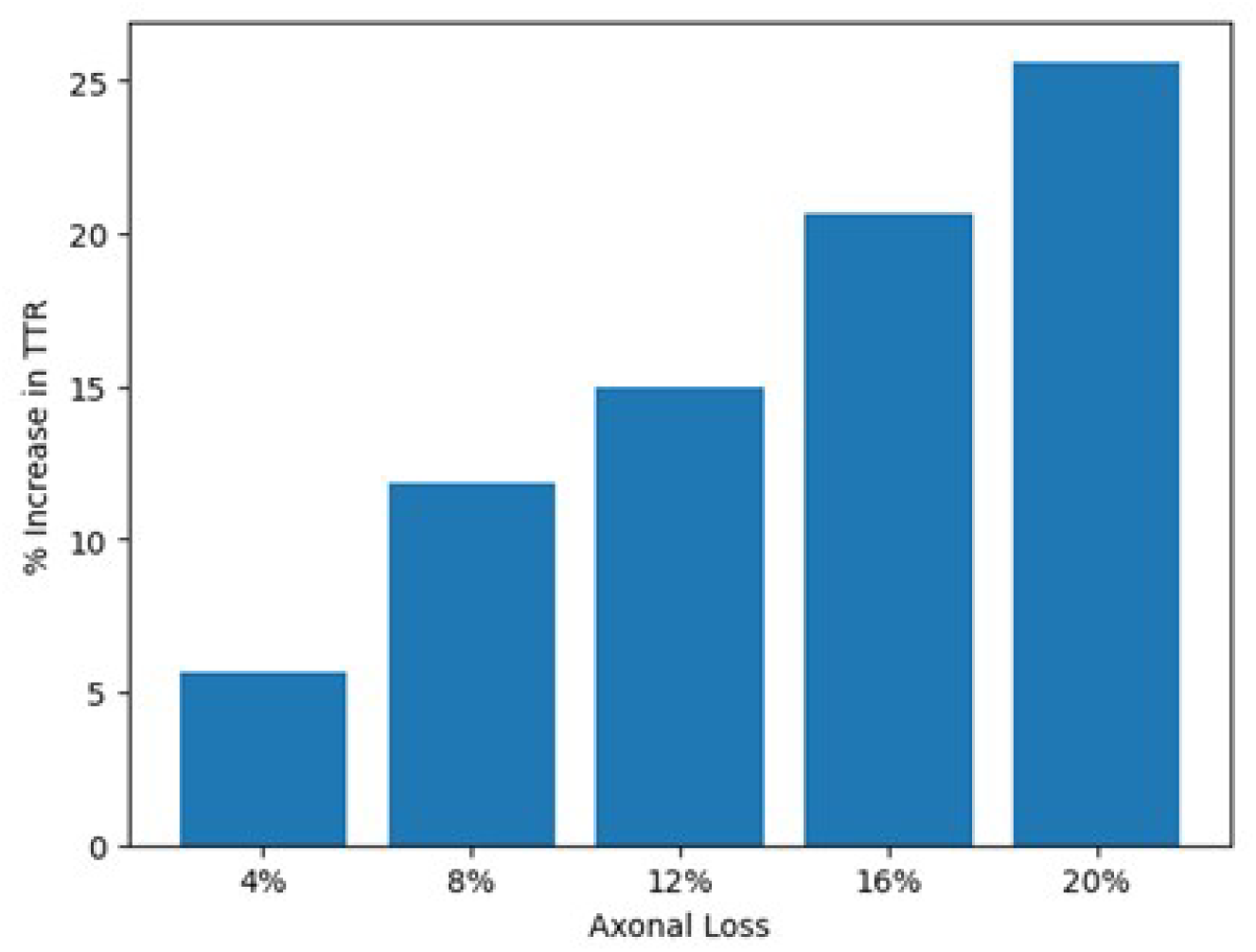
Decreased axon number increased the TTR.

### 4) Neuronal loss paradoxically decreased TTR due to memory loss which was compensated for by increased feedback

Neuronal loss is a prominent pathological feature of dementia. There is estimated to be less than 10% neuronal loss with aging, and greater than 20% neuronal loss with cortical dementias such as Alzheimer’s disease (Andrade-Moraes et al., 2013; Castelli et al., 2019; West et al., 1994; Arendt et al., 2015). We explored the effects of neuronal loss up to 20% (Figure 6). In the model, neuronal ablation in an age-associated range of up to 10% did not affect TTR (Figure 6a).

**Figure 6.**
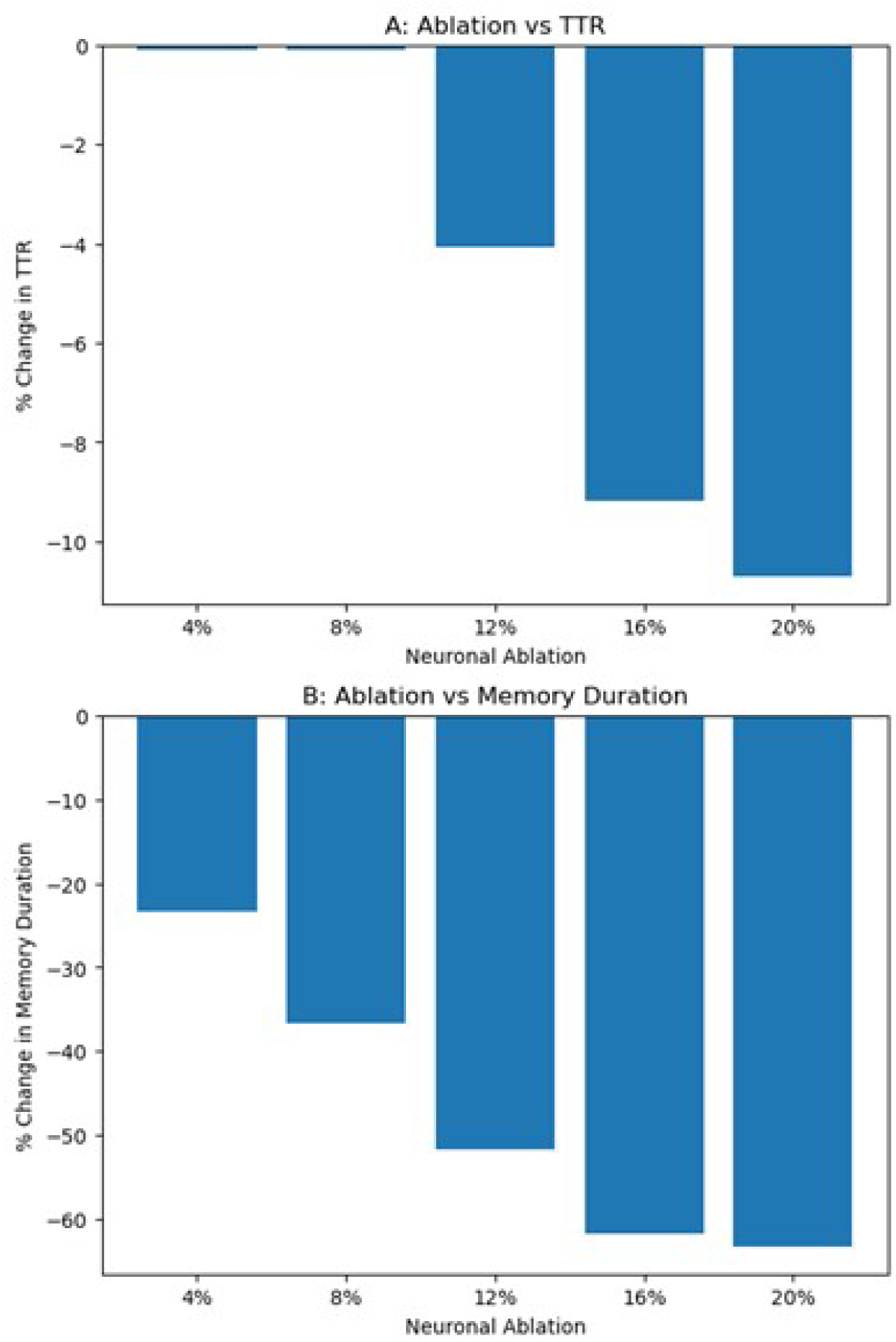
Neuronal ablation decreased both the TTR (a) and memory duration (b).

With neuronal loss above 10%, TTR paradoxically *decreased*, suggesting that the damage would lead to faster response times. Exploration of this paradoxical result demonstrated that the TTR decrease was associated with a reduction in memory duration of the current representation prior to subsequent replacement by a new representation. After the ablation, the individual memory was only partially represented by the ensemble since some of the neurons of its ensemble were absent. The memory was tenuous and could be quickly replaced by a new representation. The decrease in TTR was associated with a much larger reduction in memory duration (Figure 6b). Neuronal ablation severely impaired the recurrent circuits involved in working memory, which counteracted TTR slowing. By reducing the hold of the prior memory, the new representation could be instantiated more quickly.

Reduced feedback from reduced neuron numbers was invoked to explain the paradoxical result of Fig. 6. Feedback within the ensemble is the essence of working memory in this class of model and enjoys greater neurobiological support than do other classes of model (Murray et al., 2017; Compte et al., 2000; Wang, 2001; Lisman et al., 1998). We therefore further explored the role of feedback strength (Figure 7). As predicted, decreased feedback produced a decrease in TTR (Figure 7a). Similarly, increased feedback increased TTR. The correlate was the decrease in memory duration with reduced feedback and its increase with increased feedback (Figure 7b). As the feedback increased, the greater recurrent connectivity within the ensemble preserved the duration of working memory of the old representation, which delayed its replacement by a new representation and, therefore, increased TTR. Conversely, decreased recurrent connectivity led to tenuous memory of the old representation which was quickly replaced and decreased TTR.

**Figure 7.**
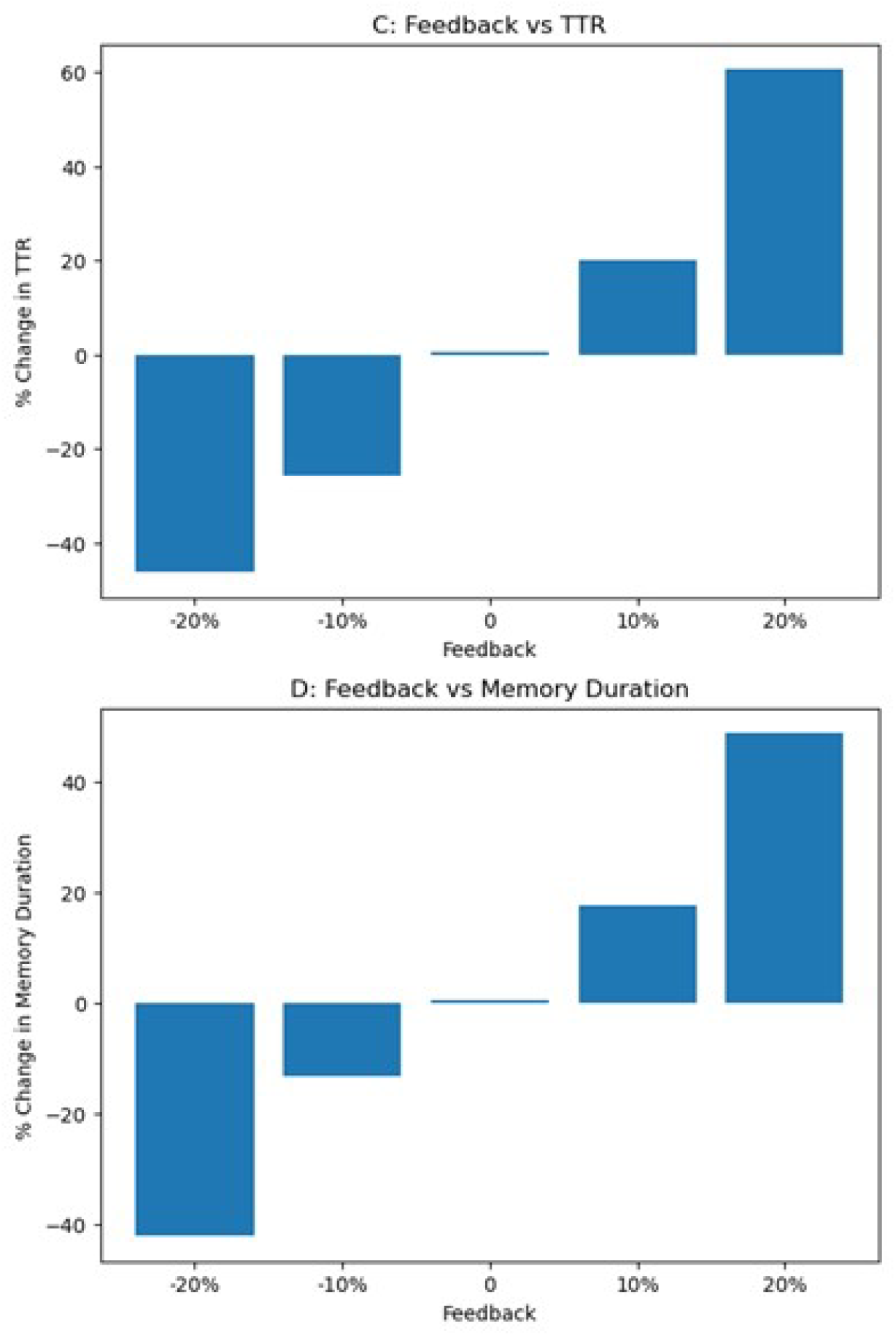
Increased feedback increased both the TTR (a) and memory duration (b).

## Discussion

Our simulations demonstrate the varied effects on part one (time to registration (TTR)) of stage one (color identification) of the Stroop test, a primary sensory task that generalizes to daily visual life. We chose the Stroop test as a framework here as it is a standard cognitive test widely used in assessing cognitive dysfunction. We assessed a set of specific pathophysiological factors that are likely to be seen singly or in combination due to particular pathologies: 1) input noise, a proxy for the signal degradation that occurs due to sensorimotor loss across modalities with aging; 2) axonal degeneration, seen as reduction in white matter noted on MRI and by pathology (Salvadores et al., 2017; Castelli et al., 2019); and 3) neuronal loss, a well-documented effect of aging (Andrade-Moraes et al., 2013; Castelli et al., 2019).

Our major finding was the increased delay consequent to most of our manipulations due to slowing of *pattern completion*. Pattern completion is a standard emergent measure of recurrent network performance (Hopfield, 1982). Incomplete input is compensated by the action of intrinsic neurons that activate each other to complete known patterns. This completing process takes time. The combination of pathological factors in the compromised individual will be cumulative in delaying signal recognition.

Pattern completion requires both the removal of anomalous activity and re-aggregating to register the correct color, requiring simultaneous processes of suppressing the anomalies and augmenting concordant information. The anomalies are due to extrinsic factors: signal noise and transmission loss. At the same time, intrinsic factors to provide pattern completion are impaired by intrinsic cortical slowing, cell loss, and transmission loss (white matter loss). As the signal-to-noise ratio decreases, more noise must be filtered out. Reconstruction of the signal requires the recruitment of additional neurons across ensembles. The steps of noise rejection, ensemble recruitment, inter-ensemble communication, and correct signal recognition are time-consuming and processing intensive, leading to cognitive delay. As sensory loss is often comorbid between sensory domains, information processing delays due to difficulty in signal perception may compound and result in considerable cognitive slowing.

Axonal loss leads to intrinsic deficits in pattern completion as the transmission of color to neurons is incomplete. When there is partial input because of axonal loss, the cortex must act as an auto-associator to gradually and recurrently activate neurons which were not directly excited due to limited input; a process which is constrained by the relatively slow time constant of recurrent excitatory synapses. The delay in excitation from cortical auto-associator activity creates a delay in signal transmission and information processing. In contrast, in the absence of axonal loss, the cortex has less need to function as an autoassociator and cognitive processing speed is preserved.

Neuronal loss causes an intrinsic deficit to pattern completion via the absence of some units of each ensemble responsible for identifying and maintaining input for particular colors. Our model suggests that neuronal loss would contribute more to memory loss than to cognitive slowing. Neurons involved in recurrent connectivity are particularly vulnerable to age-related death, which alters the firing directionality within cortical processing units. This disruption affects the activity between spatially-tuned neurons involved in recurrent activity and memory maintenance (Wang et al., 2011; Morrison and Hof, 1997). Both the reduction in neuron number, which decreases overall firing activity, and alterations in circuit connectivity between remaining neurons may contribute to reduced working memory. Unlike extrinsic deficits from signal loss, which can be compensated via cortical processing in the existing ensembles, loss of ensemble neurons requires large increases in synaptic feedback strength.

Our results suggest that both the primary degenerative changes and the compensation both play a role in contributing to reduced ability in a serial recognition task. We propose that physiological feedback is increased to preserve memory at the cost of further cognitive slowing as a way of correcting intrinsic incompleteness. The connections within an ensemble which establish recurrent excitatory activity are essential in maintaining working memory and are disrupted by neuronal loss. Increased feedback strengthens the excitatory activity which preserves working memory. Subsequently, the ensembles are made to recover memory function at the expense of further slowing - a cortical “sacrifice.” Somewhat paradoxically, this increased feedback produced a form of augmented memory, as particular color patterns were then held on to longer and were slow to be released with presentation of another color; a simple form of the perseveration seen in Wisconsin card sort and similar tests.

### Biological correlates

This feedback control may occur by adjustments in synaptic or intrinsic factors mediated by metabotropic glutamate receptors (mGluR), which have been shown to be modulated based on behavioral demands and affect cognitive performance (Sherman, 2014; Ménard and Quirion 2012).

Adjustments to the level of acetylcholine may be another biological mechanism to control the strength of recurrent excitation. Dysfunctional modulation of the gain of recurrent excitation could contribute to the disabling fluctuations in cognition that occur in dementia with Lewy bodies (Gomperts, 2016), which is linked to cholinergic dysfunction in both post-mortem (Schmeichel et al., 2008) and in vivo studies (Klein et al., 2010). Cholinesterase and butyrylcholinesterase inhibitors have been posited to have independent roles in its therapy (Kandiah et al., 2017). Palma and colleagues showed how acetylcholine could modulate the strength of recurrent excitatory connections by affecting after-hyperpolarization currents (Palma et al. 2012). Our model suggests hypotheses about cholinergic function and cognitive speed that could be tested by administering cholinesterase inhibitors in drug naïve patients and control subjects.

### Limitations of the model

The simplicity of our model of cognitive slowing is both a strength and weakness. More complex models using integrate-and-fire neurons of classic neuropsychological tasks that slow with aging may propose further factors which affect cognitive slowing and memory loss. The model identified factors responsible for cognitive slowing but lacks the multiscale molecular, synaptic and cellular factors that underlie those factors. It would be desirable to incorporate additonal changes associated with aging, such as dopamine loss in basal ganglia function.

### Prediction

The ultimate usefulness of computational models of cognition is whether they suggest empirical studies that would not be thought of without the model and whether new data collected in those studies move the field forward. We believe our model suggests several such studies. This study provides novel hypotheses about cognitive slowing that can be evaluated in neuropsychological studies correlated with imaging measures of axon integrity and cortical thinning. As above, we predict double-dissociations relating to gray versus white matter damage with respect to duration of cognitive over-holding (gray matter) compared to predominant slowing (white matter).

## Notes

Original Study. The authors have no conflicts of interest to declare. The authors are not US government employees. This research received no specific grant from any funding agency in the public, commercial, or not-for-profit sectors.

### Competing Interest Statement

The authors have declared no competing interest.

